# Climatic niche predicts the landscape structure of locally adaptive standing genetic variation

**DOI:** 10.1101/817411

**Authors:** Vikram E. Chhatre, Karl C. Fetter, Andrew V. Gougherty, Matthew C. Fitzpatrick, Raju Y. Soolanayakanahally, Ronald S. Zalesny, Stephen R. Keller

**Author notes:** Department of Biological Sciences, University of Calgary, Calgary, AB, Canada T2N 1N4, Canada. Department of Botany, University of British Columbia, Vancouver, BC, Canada V6T 1Z4.

## Abstract

Within a species’ range, intraspecific diversity in the form of adaptive standing genetic variation (SGV) may be non-randomly clustered into different geographic regions, reflecting the combined effects of historical range movements and spatially-varying natural selection. As a consequence of a patchy distribution of adaptive SGV, populations in different parts of the range are likely to vary in their capacity to respond to changing selection pressures, especially long-lived sessile organisms like forest trees. However, the spatial distribution of adaptive SGV across the landscape is rarely considered when predicting species responses to environmental change. Here, we use a landscape genomics approach to estimate the distribution of adaptive SGV along spatial gradients reflecting the expansion history and contemporary climatic niche of balsam poplar, *Populus balsamifera* (Salicaceae), a widely distributed forest tree with a transcontinental distribution in North America. By scanning the genome for signatures of spatially varying local adaptation, we estimated how adaptive SGV has been shaped by geographic distance from the rear range edge (expansion history) versus proximity to the current center of the climatic niche (environmental selection). We found that adaptive SGV was strongly structured by the current climatic niche, with surprisingly little importance attributable to historical effects such as migration out of southern refugia. As expected, the effect of the climatic niche on SGV was strong for genes whose expression is responsive to abiotic stress (drought), although genes upregulated under biotic (wounding) stress also contained SGV that followed climatic and latitudinal gradients. The latter result could reflect parallel selection pressures, or co-regulation of functional pathways involved in both abiotic and biotic stress responses. Our study in balsam poplar suggests that clustering of locally adaptive SGV within ranges primarily reflects spatial proximity within the contemporary climatic niche – an important consideration for the design of effective strategies for biodiversity conservation and avoidance of maladaptation under climate change.

## Introduction

Adaptive standing genetic variation (SGV) is of fundamental importance to the response of populations to environmental change. R.A. Fisher^1^ first identified that the response to selection is proportional to the genetic variance in fitness, but also assumed that most populations exist at or near their current fitness optima such that selection primarily acts on new mutations as opposed to SGV^2^. However, several processes are known to maintain SGV over broader temporal and spatial scales. For example, fluctuating or balancing selection may favor the long-term persistence of multiple alleles within populations, and this has been particularly well documented for genes associated with mating type^3,4^ or immunity^5–7^. Similarly, spatial population structure and local selection can maintain adaptive SGV at range-wide scales, for example due to polygenic selection shifting adaptive allele frequencies along environmental gradients^8^ or when hard or soft selective sweeps act locally within different populations^9^.

For many species, the current spatial distribution of adaptive SGV will in part reflect historical events such as range contractions and expansions in response to past glaciation, as well as effects of environmental selection during or since these events^10–13^. Hampe & Petit^14^ argued that populations distributed along the rear edge of a species’ range may be closely descended from refugial populations, and hence are predicted to be older and harbor higher levels of SGV compared to younger populations in areas of recent expansion^14,15^. Reduced SGV with distance from the rear range edge is also a predicted consequence of serial founder effects during range expansion^16^, which can lower adaptive potential with distance from rear edge source populations^17^. Moreover, rear edge populations at lower latitudes may be subject to increased levels of abiotic stress, hybridization with congeners, or experience higher disease or pathogen incidence compared to populations in the core of the species’ range^14,15,18–20^, suggesting that the strength or nature of selection may vary with proximity to the rear range edge^21,22^. However, the effect of range limits on adaptive genetic variation remains poorly understood^13,23^, and only a handful of experimental studies have quantified how SGV under selection varies along a continuum from the core to the edge of a species’ range (e.g.^24–28^).

While proximity to the rear range edge may predict if SGV for ancestral polymorphism has been reduced by colonization bottlenecks, it may not adequately capture how SGV has been shaped by spatially-varying selection after expansion^29^. In this context, populations inhabiting ecologically marginal environments are of particular interest^13,30^. Marginal environments may impose intense selection pressures on populations to locally adapt, but such adaptation may be constrained by demographic effects of genetic drift^23,31^ or the immigration of maladapted alleles from more abundant populations elsewhere in the range^32^. On the other hand, gene flow of beneficial alleles may facilitate adaptation to marginal environments^33,34^, and could contribute substantially to the global pool of adaptive SGV present across a species’ range. Identifying local populations containing unique adaptive variation relative to the rest of the range has gained increased importance in the context of contemporary global change^22^, as efforts are needed to identify and conserve adaptive germplasm *in situ* for crop breeding^35^, or to forecast the adaptive response of natural populations to climate change^36,37^.

North American tree species provide excellent study systems for addressing the effects of range context on locally adaptive SGV, as their expansion history into formerly glaciated regions is well documented in the fossil pollen record, and is known to have tracked the changing climate during the Holocene^38,39^. Due to their long individual lifespans, frequently spanning hundreds of years, trees occur in populations characterized by overlapping generations in which SGV for fitness traits can be maintained by demographic or selective processes over eons^40^. Furthermore, many tree species show strong clines across their ranges for functional traits and adaptive allele frequencies that occur across complex spatial gradients of expansion history and climatic environment^27,41–45^. This makes forest trees important systems in which to understand the spatial landscape of adaptive variation^46^, and the roles of history and selection in shaping where SGV is concentrated within the range^39^.

In this study, we report a landscape genomics analysis investigating the importance of range context for the distribution of locally adaptive genomic variation in the widely distributed boreal tree, *Populus balsamifera* L. (Salicaceae). The occurrence of *P. balsamifera* is spread across 30 degrees of latitude and over 100 degrees of longitude^47^, and as such populations span a massive range in climate and growing season length^48^. Previous genetic studies place the likely refugial location of *P. balsamifera* during the last glacial maximum (LGM) in the central Rocky Mountains^49,50^, where *P. balsamifera* currently exists in relatively isolated populations along its southern range edge^19,48^. The current center of abundance of the species lies to the north of these rear edge populations across central Canada, where population genetic and species distribution models suggest the presence of a large effective population size (*N*_*e*_) with high historic levels of gene flow to the northern and eastern peripheries of the range^49^. *Populus balsamifera* is also known to exhibit strong clinal variation for phenology and ecophysiological traits^51–53^ and for nucleotide diversity and divergence in flowering time genes, suggesting pervasive local adaptation to climate and photoperiod across its range^43,54^.

Here, we use a combination of approaches to scan the genome for signals of local adaptation, conditioned on the spatial pattern of neutral population structure, and show that adaptive SGV varies strongly with proximity to the species’ core climatic niche, with weaker effects of distance from the rear range edge. The predominant influence of the contemporary climatic niche shaping adaptive SGV is largely robust across genes involved in defense vs. drought stress, suggesting similar processes have shaped variation in genomic pathways involved in response to biotic and abiotic stress. Our landscape genomics approach gives spatially-explicit insights into where and why SGV is clustered within a species’ range, and serves as a guide for developing strategies aimed at identifying and conserving areas of unique genetic diversity and mitigating loss of adaptation under environmental change.

## Results

Our GBS libraries for 508 individuals generated 3.9 billion single-end reads passing filter, of which 1.35 billion passed QC in the GBS pipeline as “good, barcoded reads” *sensu* Glaubitz et al.^55^, consisting of a perfect match to one of the barcodes and no ambiguous base calls. Reads passing QC clustered into 3,755,656 sequence tags that mapped uniquely to 220.9 Mb of the *P. trichocarpa* reference genome v3.0. Variant calling produced 1,107,538 sites which, after filtering, resulted in 167,324 biallelic SNPs for inference of population structure and selection, with a median (across individuals) per-site depth of 21.8X.

### Population Structure

Inference of population structure via discriminant analysis of principal components (DAPC) revealed support for 3 to 4 genetically distinct clusters, with mean BIC over the 1000 iterations minimized at *K*=3 (Fig. S1A, B). Clustering assignments at *K*=3 separated individuals into a northwestern cluster in Alberta and British Columbia, a central cluster throughout the range core from Saskatchewan to Ontario including the Great Lakes region, and an Atlantic Canada cluster along the eastern coast (Fig. 1C inset). In about 10% of the iterations, DAPC favored a model of *K*=4 that showed a slightly higher BIC averaged across iterations compared to *K*=3. At *K*=4, the dominant solution split the range core into eastern and western clusters that formed distinct groups but were only weakly differentiated (*F*_ST_ = 0.008) (Fig. 1C).

**Figure 1.**
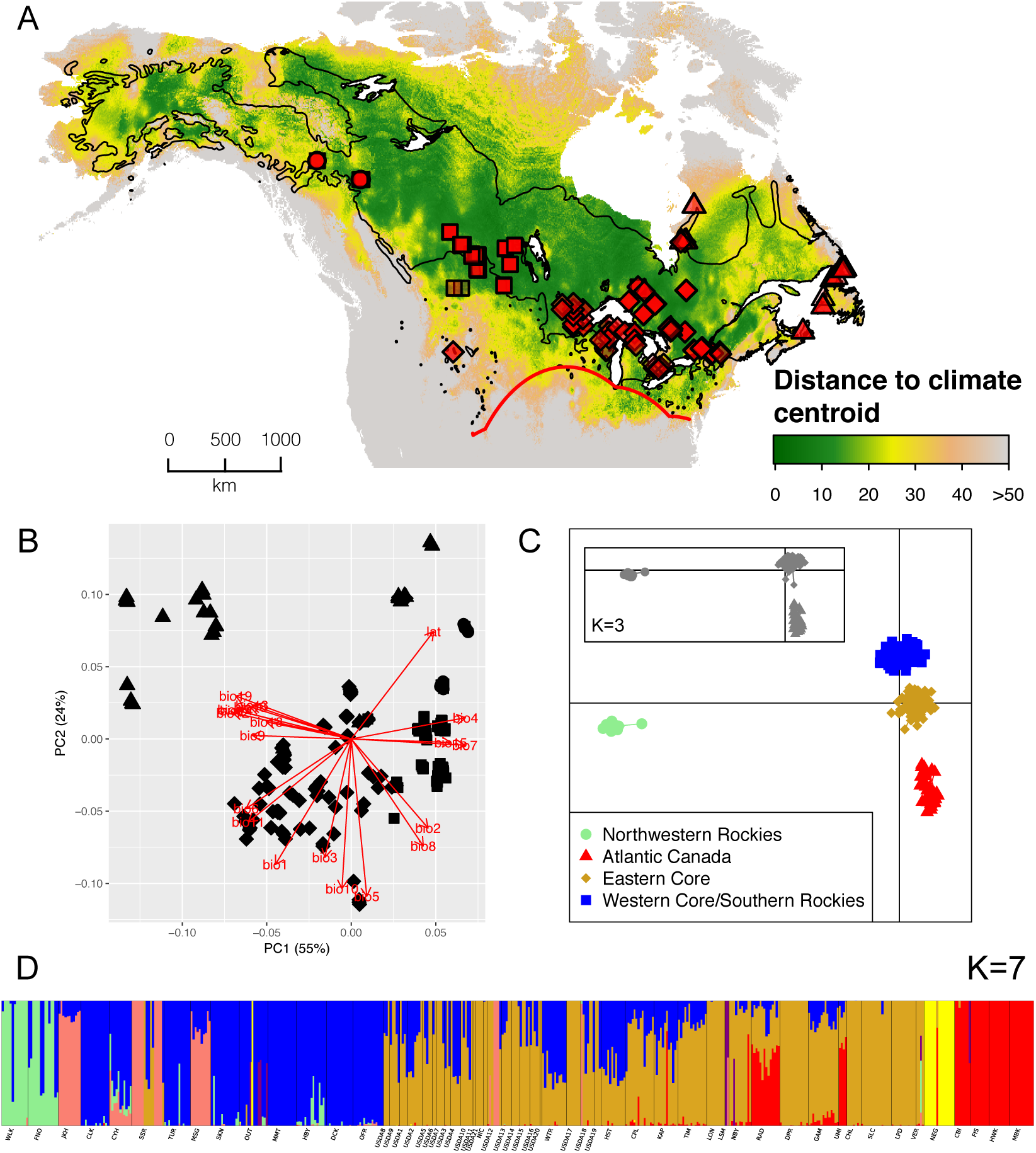
Sample collection map of the 437 individuals analyzed in selection scans. The red line represents the estimated location of the southern range edge based on occurrence records (A). The range boundary of *P. balsamifera* is denoted by the black line and is adapted from^47^. 19 BIOCLIM variables and latitude were included in a PCA and PC1 and PC2 were used in gene-environment analyses (B). Discriminant analysis of principal components (DAPC) at *K* =4 separates Northwestern Rocky Mountains from populations east of Saskatchewan along DAPC axis 1. DAPC axis 2 separates an Atlantic Canada clusters and two range core clusters. Shapes in the sample map, PCA plot correspond to the DAPC (C). ADMIXTURE clustering of all 508 individuals sequenced in the study at *K* =7, the level of K with the lowest cross validation error. Colors represent the following clusters: blue & goldenrod: range core, lightgreen: Rocky Mountain region, red: Atlantic Canada, yellow: New Glasgow, NS, Canada with introgression from *P. deltoides*, pink: admixture from Northwestern cluster reported by Keller *et al*^49^. Individuals with less than 90% *P. balsamifera* ancestry were removed prior to performing selection scans.

The genetic clusters from DAPC aligned along major gradients of climate and growing season length, as captured by the first two principal components of the climate PCA (Fig. 1B). Large values of PC1 represented dry environments with high temperature and precipitation seasonality, and were primarily occupied by the western core and northwestern Rockies groups, while small values of PC1 (wet, low seasonality) were occupied by individuals with ancestry in the Atlantic Canada cluster. PC2 represented a north-south gradient of growing season length, with large values representing high latitude, cool sites occupied by the northwestern Rockies and a subset of the Atlantic Canada clusters, while small values of PC2 (low latitude, warm sites) were dominated by members of the eastern core cluster (Fig. 1B). Ancestry clusters were also non-randomly distributed with respect to distance from the rear range edge and proximity to the species-wide climatic niche centroid (Fig. 1A). Notably, the Atlantic Canada cluster was farthest from the climatic centroid while spanning a range of distances from the rear edge. In contrast, the western and eastern core clusters were close to the climatic centroid, with many eastern core populations also proximate to the rear range edge (Fig. 1A).

ADMIXTURE analysis largely corroborated the population structure found by DAPC, but also revealed extensive mixed ancestry among *P. balsamifera* clusters, as well as introgression with other *Populus* species (Fig. 1D). At lower levels of *K* (2–3), introgression was especially prevalent with *P. trichocarpa* along the rear range edge and in the northwestern Rockies (Fig. S2; see also Chhatre *et al*^19^). At *K*=4, a separate source of introgressed ancestry was apparent in the Atlantic Canada population of NEG, representing hybridization with *P. deltoides*^20^. Scattered introgression was also apparent at a low frequency throughout other portions of the range. At higher levels of *K* (5–6), ancestry within *P. balsamifera* separated into an Atlantic Canada cluster and a subdivided central range core split into eastern and western clusters, as observed in DAPC (Fig. 1D). Individual ancestry assignments to the eastern and western range core revealed a longitudinal cline, indicating extensive shared ancestral polymorphism and/or gene flow between these two weakly differentiated groups. At *K*=7, the best model supported by ADMIXTURE cross-validation, an additional distinct ancestry cluster was evident in the far northwestern Rockies (WLK and FNO; Fig. 1C).

### Genomic Signals of Local Adaptation

After eliminating individuals with less than 90% *P. balsamifera* ancestry (N=26 individuals; Fig. 1D), we retained 437 individuals and 129,251 SNPs to test for genomic signatures of local adaptation, with 67% of SNPs within 5 kb of an annotated gene model. Genome scans identified 397 outlier SNPs in tests for elevated population differentiation (Bayescan and *X*^*T*^ *X*) that exceeded the empirical Type I error rate based on the distribution of intergenic SNPs (*α*=0.0006) – an enrichment of more than 2.5X over neutral expectations. Most differentiation outliers were significant only in Bayescan (321/397 loci, 81%) or to a lesser extent, in *X*^*T*^ *X* (5 loci, 1%), while 71 outliers (18%) overlapped between the two differentiation tests (Fig. 2a). Gene-environment association (GEA) tests (LFMM and Bayenv2) returned 535 outliers associated with either climate PC1 or PC2, more than 1.7X greater than neutral expectations. Similar to differentiation tests, most outliers in GEA tests were sensitive to the method used to detect them, with 16 of 535 loci (3%) in the range-wide set shared between Bayenv2 and LFMM (Fig. 2a). Combining across all four genome scan approaches, we identified 814 loci within 5kb of 724 genes that showed evidence of either elevated differentiation or gene-environment association relative to the empirical neutral distribution, with most loci unique to a single analysis (85%) and only 6 loci (0.7%) shared across all four methods.

**Figure 2.**
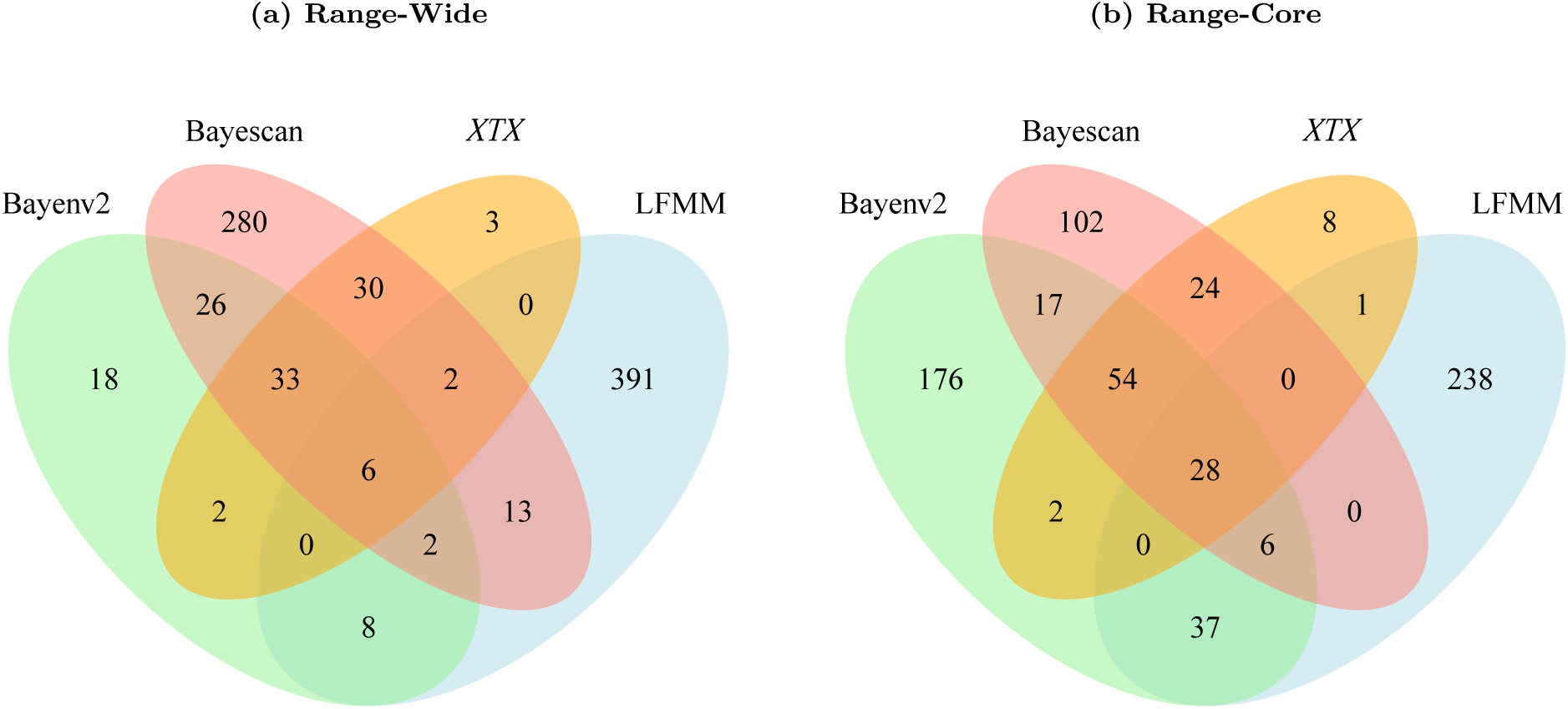
Venn diagrams showing overlap of local adaptation outliers among four selection scans using the Range-Wide and Range-Core samples. Bayenv2 and LFMM test for gene-environment associations (GEA), whereas Bayescan and *X*^*T*^ *X* test for elevated population differentiation.

Outliers unique to a single method may be false positives caused by inadequate control of background structure due to residual relatedness or departure from the assumed demographic history. Alternatively, these loci may be true positives that are not detected by the complimentary GEA or differentiation method, for example if the statistical modeling of background genetic structure over-corrects the allele frequencies and dilutes the selection signal. To explore this possibility further, we performed each selection scan again on just those populations belonging to the eastern and western range core clusters, where background population structure was minimal (Fig. 1; N=336 individuals; *F*_ST_ *<* 0.01). This greatly increased the frequency of shared outliers among differentiation methods to 44% (106/242 loci; Fig. 2b), and confirmed several prominent genomic regions of high differentiation in both the range-wide and core sets (Fig. 3). These genomic regions included a large peak encompassing ca. 168 kb on the distal arm of chromosome 3, a second prominent peak spanning ca. 369 kb on chromosome 4, and an outlier marked by a single SNP on chromosome 8 (Fig. 3).

**Figure 3.**
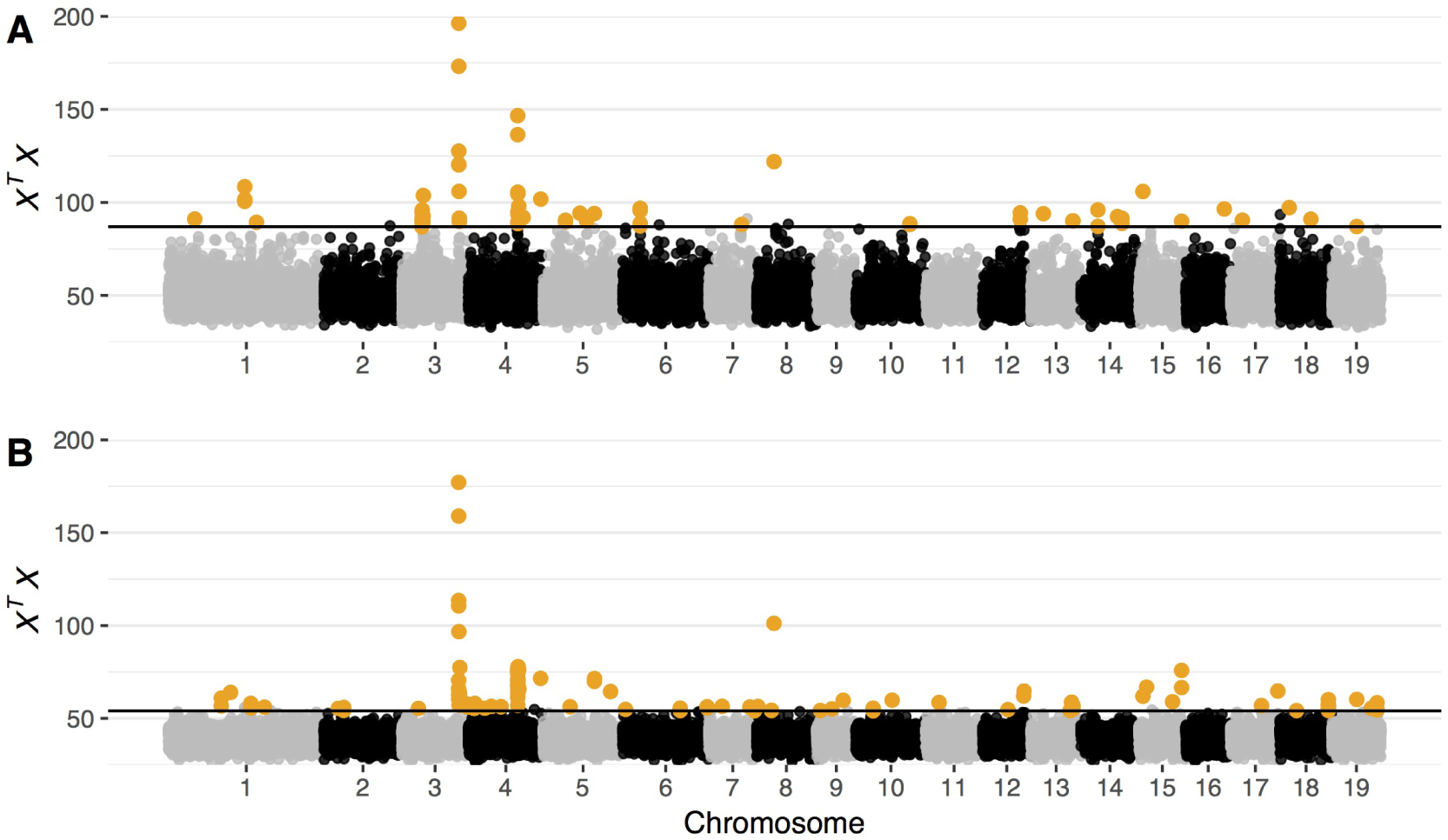
Manhattan plots of *X*^*T*^ *X* outlier scan for the set of range-wide individuals (A) and the range core individuals (B). Highlighted in each panel are outlier loci that were also identified in Bayescan; 106 loci in panel A and 71 in panel B. The horizontal black line indicates the -log_10_*P*-value significance threshold based on the empirical neutral distribution

A more modest increase in the frequency of shared outliers was evident between GEA methods performed on the range core set (13%; 71/559 loci); however, the number of outliers identified by Bayenv2 increased dramatically, suggesting the selection signal in the range-wide set likely contained false negatives due to over-correction of population structure (Fig. 2b). Genomic regions of shared outliers in the range-wide set were limited to a few scattered chromosomal regions, whereas in the range core, shared outliers were clustered into prominent peaks on chromosomes 3, 4, 8, 12, 15, and 17 (Fig. 4).

**Figure 4.**
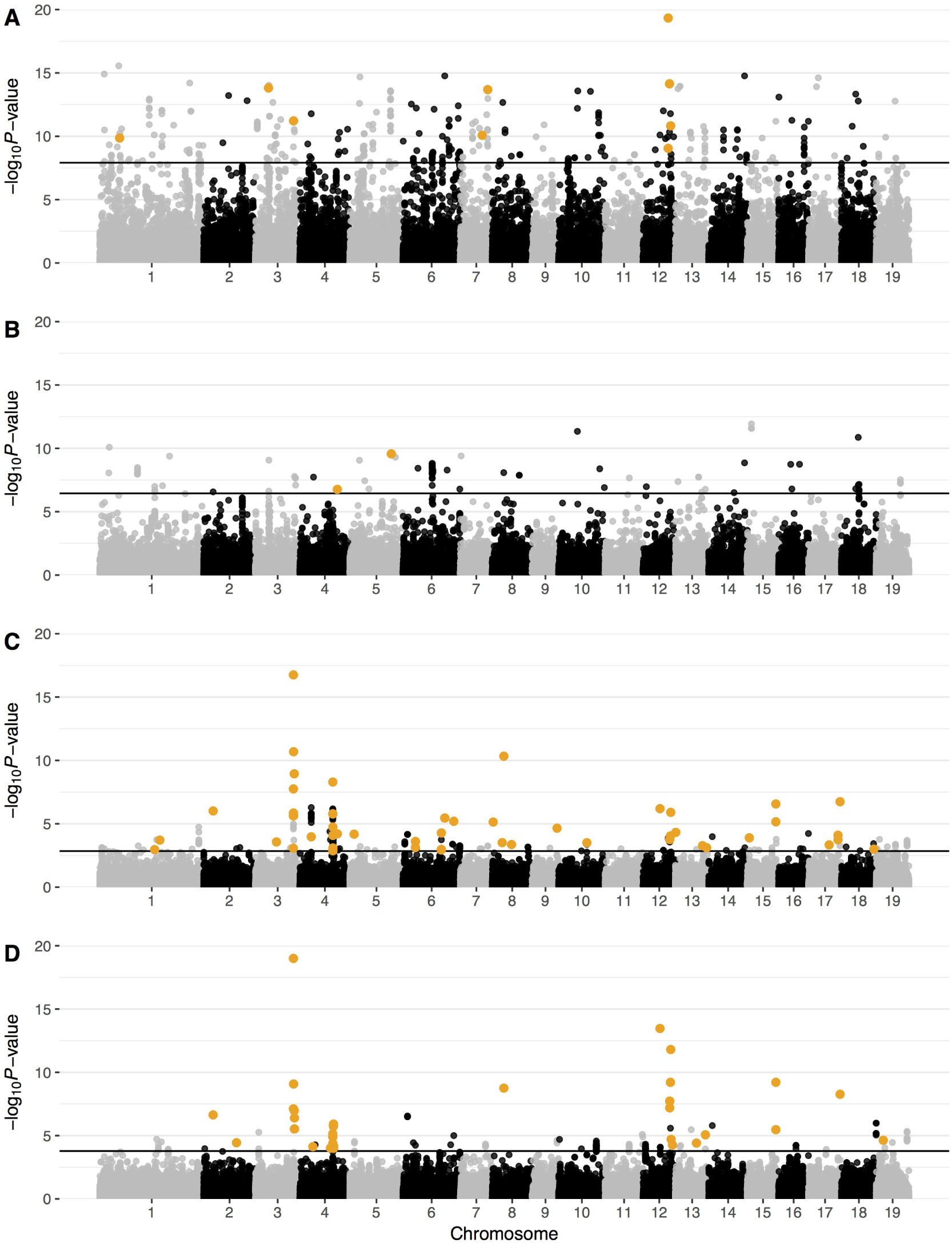
LFMM results for gene-environment association tests for the range-wide (A,B) and range core (C,D) sample sets with climate PC1 (A,C) and PC2 (B, D). Loci in orange were also significant in the Bayenv2 gene-environment test. The horizontal black line indicates the -log_10_*P*-value significance threshold based on the empirical neutral distribution

Across all four methods, inference of loci under local adaptation appeared to be sensitive to the genome scan approach (differentiation vs. GEA) and the specific method used. Despite multiple controls in place for background genetic structure, false positives are inevitably part of these differences. However, the genetic architecture of local adaptation is likely to be dominated by many loci of small effect that exhibit weak allele frequency (co)variance among populations, making it difficult to distinguish these loci from the genomic background.

Additionally, some loci likely experienced selection along portions of the climatic gradients that run parallel with the neutral genetic structure (e.g, climate PC2 and the Atlantic Canada cluster; Fig. 1). Accordingly, we retained two sets of outliers for further investigation: (1) an inclusive set of all 814 outlier loci identified by one or more selection scan methods in the range-wide analyses, and (2) only those outliers corroborated by both methods within a given genome scan approach (differentiation or GEA), resulting in a reduced set of 203 loci. The former is more likely to retain true positive cases of weakly selected loci or those spatially confounded with neutral structure at the expense of including more false positives, while the latter is less likely to contain fewer false positives but will be biased towards loci of large effect (e.g., large allele frequency differences) that are not closely aligned with neutral population structure.

### Distribution of adaptive SGV along geographic and climatic range edges

We investigated the spatial turnover in SGV by determining the relative contribution of individual populations to the adaptive variation we observed across the entire set of populations. For each population, we estimated the population adaptive index (PAI) of Bonin *et al.*^56^:

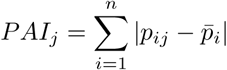

where *p*_*ij*_ is the minor allele frequency of the *i*th outlier locus in the *j* th population, and 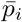 is the mean frequency across all populations. Populations with high values of PAI contribute disproportionately to the pool of adaptive allele frequencies; while populations with low PAI have allele frequencies closer to the range-wide average. To assess how this among-population component of SGV varied across the landscape, we modelled PAI as a function of distance from the rear edge and the climatic niche centroid. When considering the inclusive set of 814 selection outliers as an index of genomic beta diversity, over 75% of the variability in PAI was explained by jointly accounting for geographic distance from the rear edge and the euclidean distance from the climatic niche centroid (Table 1). Contrary to predictions based on expansion history^12,14^, PAI showed little effect of distance from the rear range edge except through an interaction with climate distance, which revealed a dramatic increase in PAI with combined distance away from both the rear edge and climatic niche centroid, consistent with divergent local selection in climatically marginal environments that are geographically located far from the current rear range edge (Fig. 5a). Driving this trend were large values of PAI for populations in Atlantic Canada that inhabited the easternmost portion of the species’ range – a region climatically distinct from the majority of the species’ niche (Fig. 1A, B; Fig. 5a). The overall trend in PAI was similar for the large effect outliers (N=203 loci), although this set of loci showed some rear edge populations exhibiting slightly elevated levels of PAI but still far below levels seen in marginal climates (Table 1; Fig. 5b).

The historic and selective processes shaping the geographic distribution of SGV may differ for adaptive variation underlying responses to abiotic vs. biotic stress^13,18^. To test this prediction, we calculated PAI separately for outlier SNPs that tagged genes showing differential expression under controlled drought (abiotic) or beetle/mechanical wounding (biotic) based on the gene expression atlas on the Populus Genome Integrative Explorer (PopGenIE)^57^. Among the 724 genes associated with the set of inclusive outliers, 105 genes were differentially expressed under drought (N=174 outlier SNPs), and 30 were differentially expressed under beetle or mechanical wounding (48 outlier SNPs). Both types of stressor showed a strong increase in PAI towards climatically marginal range edges, although the effects were more pronounced for loci associated with abiotic stress response (Table 1, Fig. 5c,d). Interestingly, PAI for biotic stress outliers increased consistently with distance from the rear edge in a trend that was not evident among drought-associated loci (Fig. 5c).

**Table 1.**
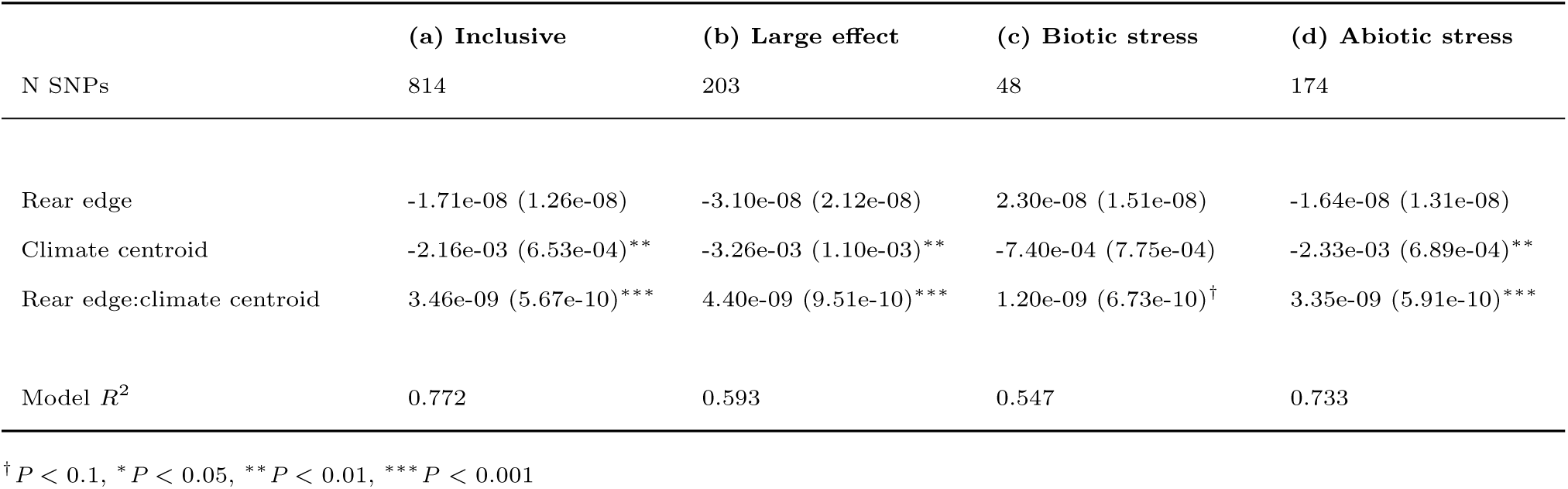
Response of population adaptive index (PAI) to distance from the rear range edge and from the climatic niche centroid. (a) The inclusive set of significant outliers identified across all four selection scan methods; (b) large effect outliers corroborated by both GEA-based tests and/or both *F*_ST_-based tests; (c,d) outliers proximate to (¡5kb) genes differentially expressed in response to (c) biotic (beetle or mechanical) or (d) abiotic (drought) stress. Values are coefficients (SE) from multiple regression.

**Figure 5.**
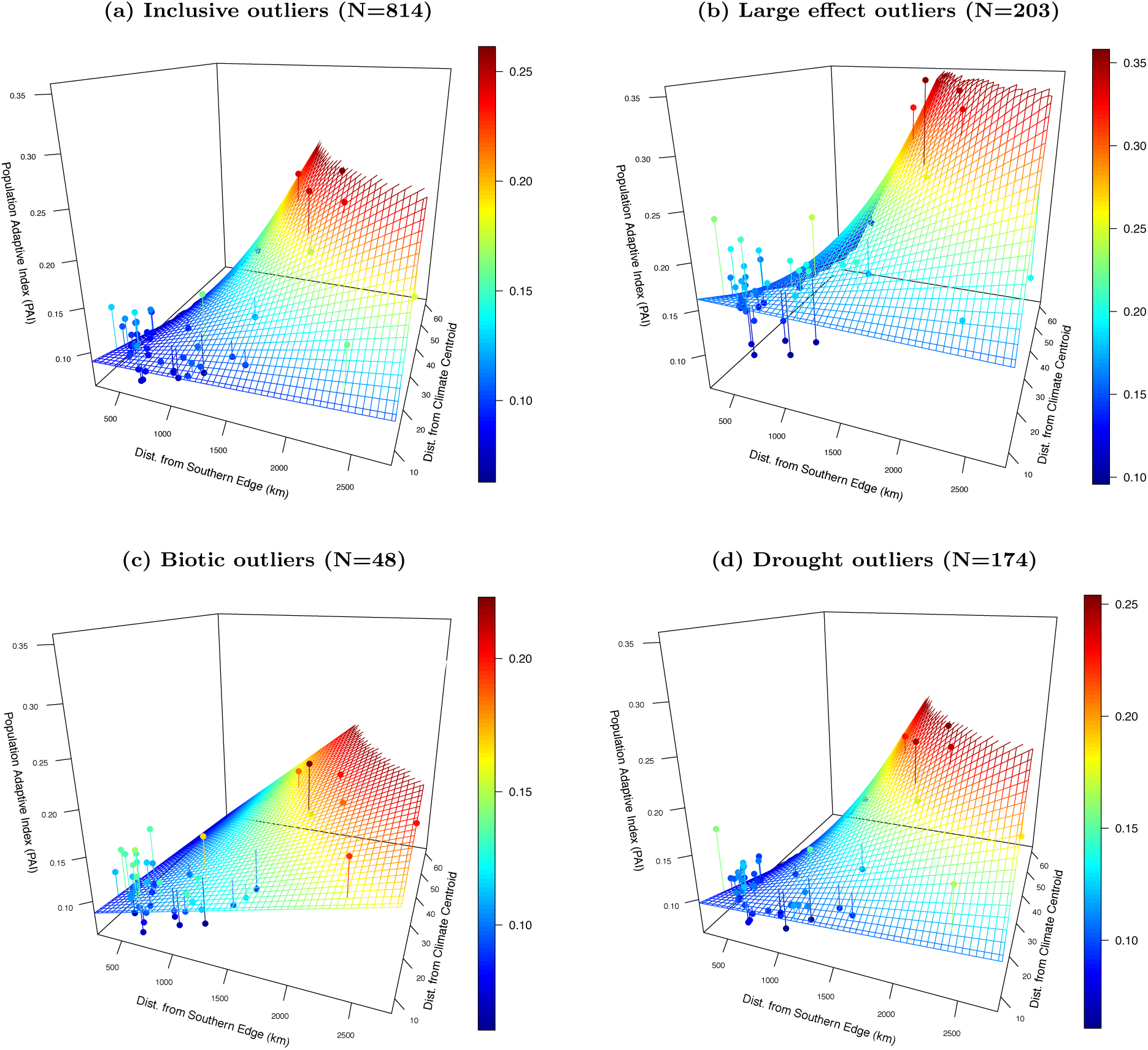
Predicted model fit surface of the relationship between population adaptive index (PAI) and two predictor distances (distance from the rear range edge and distance from the climate centroid). PAI estimates are summed over all outliers and normalized for comparison between data sets (see definitions in Table 1). Surface colors show PAI magnitude from low (blue) to high (red).

As an index of genetic beta diversity, PAI identifies populations whose allele frequencies deviate most from the sample-wide mean, and thus this estimator is inherently on the *among*-population component of SGV. To investigate the effects of range position on *within*-population SGV (i.e., alpha diversity), we estimated the variance in allele frequencies at biallelic sites (*pq* =*p*\*(1-*p*)) for each population averaged across outlier loci. The inclusive set of 814 outliers showed a highly significant interaction between rear edge and climatic distances (Table 2), with *pq* for locally-adaptive alleles highest with increasing distance from the rear edge but close to the center of the climatic niche (Fig. 6a).

**Table 2.** Response of allele frequency variance (*pq*) averaged over outlier loci to distance from the rear range edge and from the climatic niche centroid. Variable definitions and outlier groups are as described in Table 1

|  | (a) Inclusive | (b) Large effect | (c) Biotic stress | (d) Abiotic stress |
| --- | --- | --- | --- | --- |
| N SNPs | 814 | 203 | 48 | 174 |
| Rear edge | 2.02e-08 (4.10e-09)*** | 9.79e-09 (8.79e-09) | 1.41e-08 (5.46e-09)* | 2.13e-08 (4.20e-09)*** |
| Climate centroid | 5.46e-04 (2.11e-04)* | 4.17e-04 (4.55e-04) | -1.22e-04 (2.83e-04) | 3.52e-04 (2.17e-04) |
| Rear edge:climate centroid | -7.63e-10 (1.83e-10)*** | -6.67e-10 (3.95e-10)<sup>†</sup> | -2.57e-11 (2.46e-10) | -7.31e-10 (1.89e-10)** |
| Model $R^2$ | 0.386 | 0.111 | 0.388 | 0.466 |

<sup>†</sup> $P < 0.1$ , \* $P < 0.05$ , \*\* $P < 0.01$ , \*\*\* $P < 0.001$

**Figure 6.**
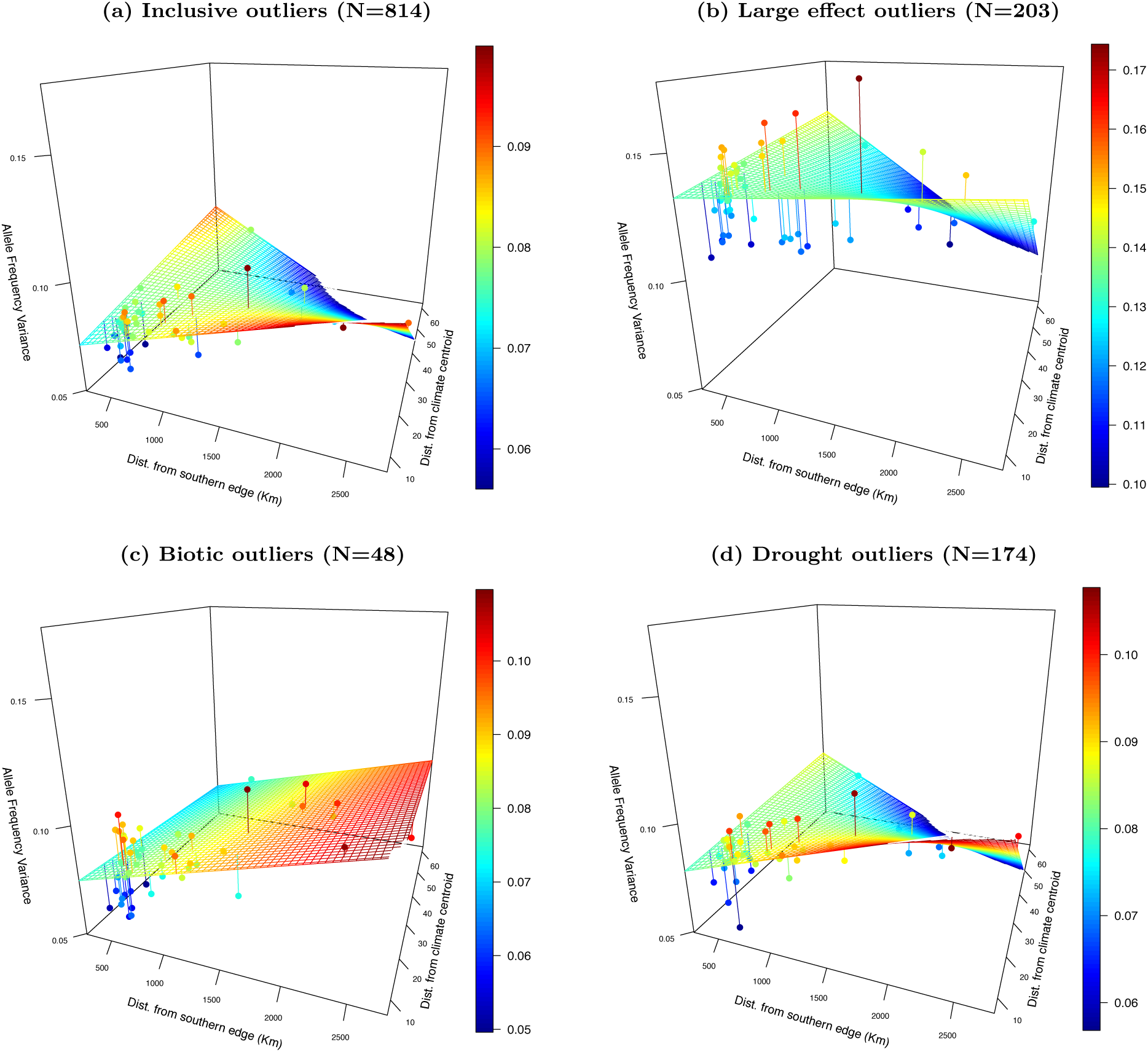
Predicted model fit surface of the relationship between allele frequency variance (*pq*) and two predictor distances (distance from the rear (southern) range edge and distance from the climate centroid). Panels (a) through (d) correspond to categories of outliers decribed in Figure 5. The z-axis is scaled to span the range of allele frequency variance observed across all four comparisons (min=0.04; max=0.17). Surface colors indicate strength of allele frequency variance from low (blue) to high (red).

For large effect outliers and those associated with drought stress, allele frequency variance was lowest on average for climatically distant populations where PAI was high, suggesting selection has depleted SGV locally (*pq*) while increasing the genetic distinctiveness (PAI) of climatically marginal populations (Fig. 6). In contrast, within-population SGV for outliers associated with biotic stress increased significantly with distance from the rear edge (Table 2), but showed no consistent effect of position along the climate gradient (Fig. 6).

The effects of distance from the rear range edge and climatic niche on components of adaptive SGV are evident when population values of *pq* and PAI are mapped onto the landscape (Fig. 7). With the exception of the biotic outliers, the overall trend showed higher within-population SGV and lower among-population SGV nearer to the geographic and climatic center of the range, per theoretical expectation of the ecological and evolutionary processes structuring range limits^58^. Two exceptions to this general trend were evident. First was the clearly elevated among-population SGV seen in the Atlantic Canada populations, and that featured prominently in all outlier sets. Second was the unusual combination of SGV associated with biotic stress response genes, in which Atlantic Canada populations harbored high levels of SGV both within- and among-population (Fig. 7).

**Figure 7.**
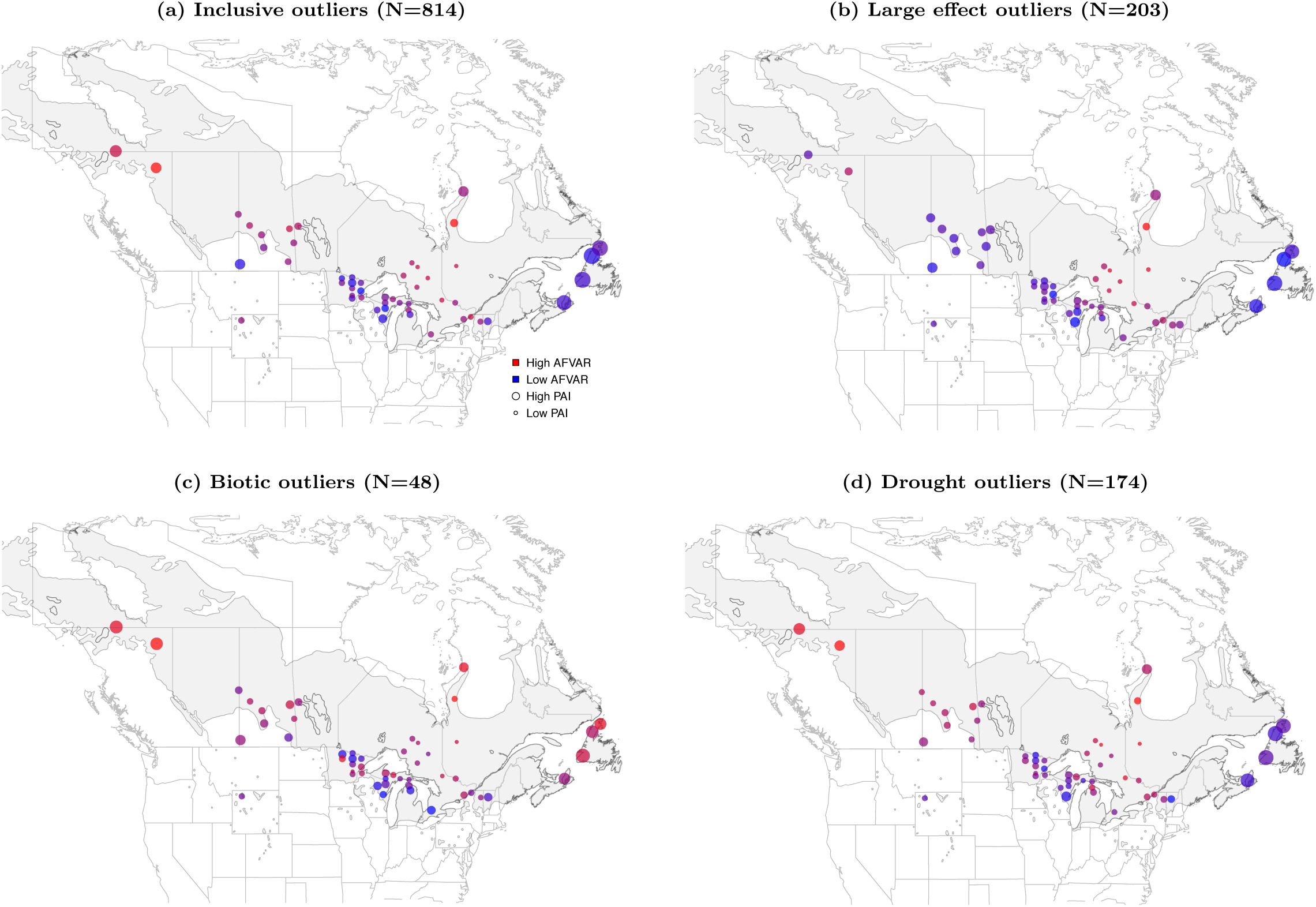
Landscape level comparison of the relationship between allele frequency variance (color: blue to red) and population adaptive index (size: small to large) among four groups of selection scan outliers. PAI point sizes are not comparable between individual subplots.

### Functional enrichment of adaptation outliers

We found 52 gene ontology (GO) terms significantly enriched among genes associated with the 814 outlier set (Supplementary Information). These included clusters of terms relating to enzyme transferase activity, biosynthesis of cell wall components such as cellulose and lipids, signal transduction and ROS activity, response to stress and/or DNA damage repair, and transport across cell membranes (Fig. 8). Genes associated with the 203 outlier set showed significant enrichment for 28 GO terms including many similar functions as the 814 outliers set, but with the addition of several terms associated with vessicle transport within and across cell membranes, and response to foreign bodies (Supplementary File S1).

**Figure 8.**
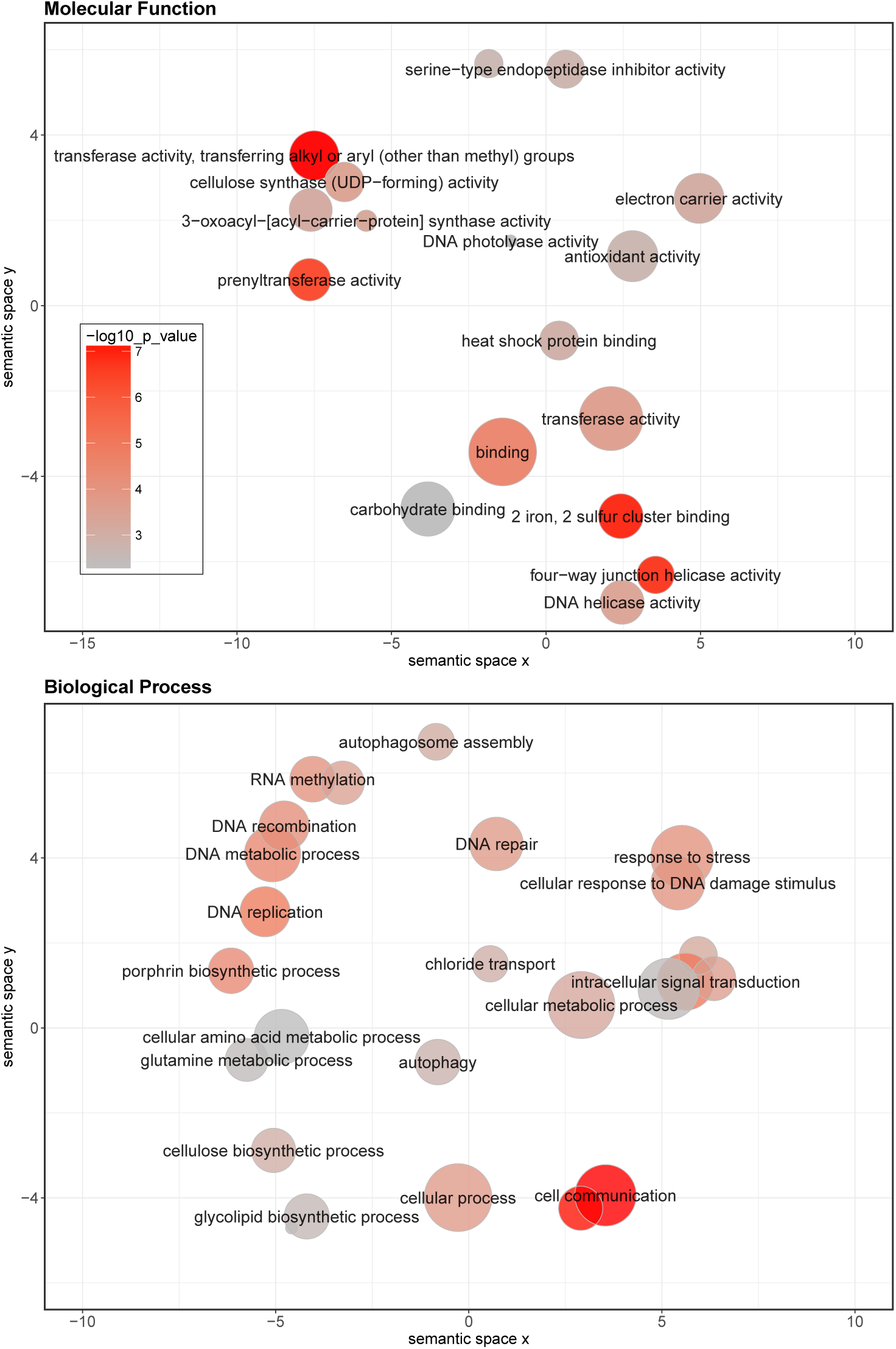
Redundancy clustering of functionally enriched GO terms associated with the 814 loci significant in the range-wide local adaptation scans. Color indicates significance of the enrichment for each GO term, and symbol size is proportional to the frequency of each GO term in the *P. trichocarpa* v3.0 genome annotation

Several of the enriched GO terms were associated with candidate genes tagged by SNPs that were selection outliers in all four genome scan methods in the range-wide and core sets (Table 3) These included candidate genes within the selected region on chromsome 3: the flavonoid biosynthesis gene CHALCONE SYNTHASE, CELLULOSE SYNTHASE-LIKE D1.1, and two FERRODOXIN genes. Chromosome 12 also contained candidate genes with enriched GO terms associated with DNA repair: an ATPase-like family protein (Potri.012G096300), RESPIRATORY BURST OXIDASE HOMOLOG C, a Tyrosyl-DNA phosphodiesterase, and a UvrD DNA helicase. Chromosomes 15 and 17 each contained a single gene with enriched GO terms: the MADS-box transcription factor TRANSPARENT TESTA 16, and the stress-responsive molecular chaperone HEAT SHOCK PROTEIN 90-1 (Table 3).

**Table 3.**
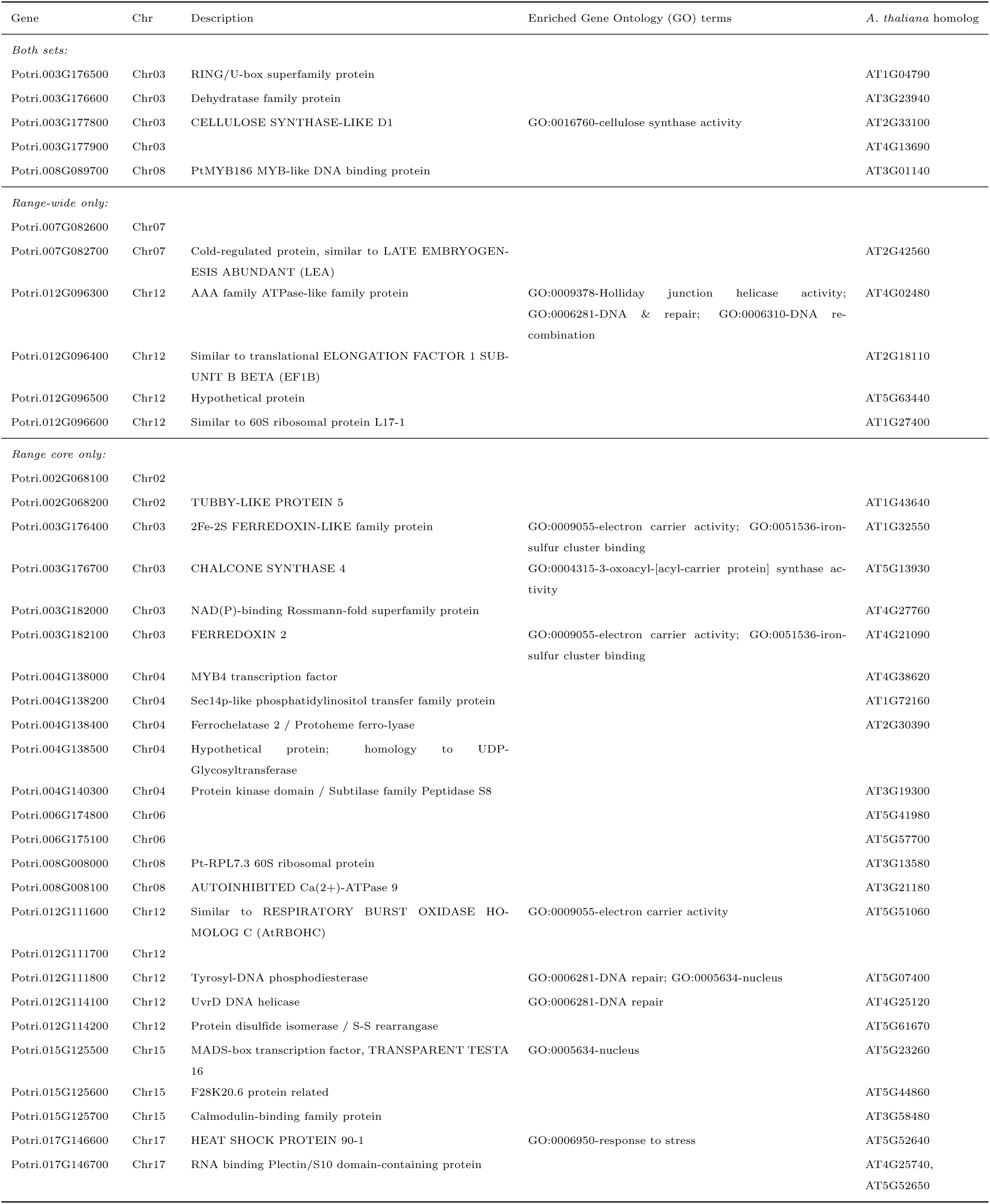
Candidate genes for local adaptation in range-wide and range core datasets. Annotated genes are located within 5kb of outlier SNPs identifed by all four selection scan methods.

## Discussion

The current period of rapid global climate change has created an immediate need to understand the genetic basis of climate adaptation in natural populations. For long-lived sessile organisms such as forest trees, adaptation to ongoing climate change will depend in large part on the availability of adaptive standing genetic variation (SGV) within the species, as well as the distribution of SGV across a potentially expansive landscape. Numerous studies have shown the post-glacial biogeography of forest trees shaped SGV over millennia, involving complex cycles of range contraction and expansion, and also local adaptation along climatic gradients^10,14,40,41^. But while forest trees have become important models for understanding the genetic basis of climate adaptation at landscape scales^46,59,60^, few studies have explicitly addressed how adaptive SGV is partitioned along spatial gradients reflecting expansion history and current climatic niche^13^.

Our approach fills an important gap in identifying geographic regions where adaptive genomic diversity is concentrated within the range, while also revealing the relative importance of historic and selective processes shaping the current distribution of SGV^13,29,61,62^. Our findings in a widespread forest tree species suggest that adaptive SGV clusters non-randomly along geographic and climatic gradients, with decreasing alpha diversity and increasing compositional turnover (i.e., beta diversity) with distance away from the center of the current climatic niche (Fig. 5 and 6). Contrary to expectations of loss of diversity due to founder effects during expansion from refugia, we observed relatively low SGV in low latitude populations^14^. These findings strongly suggest that regionally common SGV is maintained in the climatic center of the range, while climatically marginal populations are unique in harboring locally reduced but regionally rare adaptive variation.

In many ways, our findings mirror the Centre-Periphery Hypothesis (CPH), that predicts within-population genetic diversity should be highest near the abundant center of the range, while population differentiation should increase towards the range periphery where environments become less optimal and abundance drops^13,58^. Gougherty et al.^63^ also found support for the CPH in poplar, using a larger range-wide sample of populations and focusing exclusively on the alpha diversity of putatively neutral SNPs sampled randomly from the genomic background. This suggests that both neutral and adaptive genomic diversity follows the predictions of the CPH because both are jointly influenced by *N* e. That is, we might expect higher alpha and lower beta diversity across the genome where the region of largest *N* e (which presumably reflects the demographic abundance of the species) is centered with the species’ environmental niche. This assumes the major selective influences shaping range limits are climatic in origin, thus aligning climate-adaptive variation with spatial locations of demographic abundance; however, this expectation might break down for genes under strong selection for functional responses not directly related to climate^13^. The majority of the genomic outliers for adaptation studied here fit this prediction, with high alpha diversity (*pq*) and low beta diversity (PAI) near the geographic and climatic center of the range, consistent with the CPH.

One notable exception occurred for outliers associated with response to biotic stress, which showed alpha diversity increased monotonically away from the rear edge, while beta diversity also increased with distance from the climatic niche (Fig. 6). Thus, genes associated with defense response appear to harbor regionally high levels of diversity in the same climatically marginal region of Atlantic Canada, while also maintaining high levels of within-population polymorphism for these genes. High levels of within-population polymorphism that are also regionally unique could be caused by several evolutionary processes, such as a history of spatial or temporally varying balancing selection, or selection-driven introgression from congeners. Both processees would be expected to lead to high levels of polymorphism within populations that may also be unique to the region where the selection or introgression is occurring. Regardless of the mechanism, the high alpha and beta diversity for biotic response in Atlantic Canada suggests the selective history on defense-related genes may be unique in this portion of the range.

The Atlantic Canada populations (in our study, populations CBI, HWK, MBK & FIS) occupy a maritime climate quite different from the rest of the range, characterized by cool temperatures and high precipitation amounts with low seasonality^48^. We previously showed that balsam poplar in this region were divergent from the rest of the range for a suite of phenology and ecophysiological traits associated with climate adaptation, and that this phenotypic divergence was in excess of that predicted by background genetic structure, suggesting a unique history of local selection in Atlantic Canada^52^. In a separate study of *P. balsamifera*, Meirmans *et al*^64^ similarly reported a genetically distinct Atlantic Canada group that showed little admixture with the nearby range core ancestry group to its west, which they suggested was likely rooted in the local adaptation of the Atlantic Canada group to the unique maritime climate. Consistent with this proposition, the PAI for most of the local adaptation outliers in our study peaked in the Atlantic Canada group, whereas within-population allele frequency variances were lowest in this group, suggesting local selection driving SGV adaptive under these climate conditions towards fixation (Fig 5b, d & 6b, d).

It is important to note that landscape genomic studies of local adaptation invariably capture other selective gradients that are geographically aligned with climate; thus, what we infer as genes under climate-driven selection based on associations with temperature or precipitation gradients may actually reflect other unknown causal agents of selection. In our range-wide analysis, we found outlier genomic regions were significantly enriched for a variety of functions, including oxidoreductase activity, response to stress/DNA damage repair, cell signaling/transport, metal (iron) binding, and biosynthesis of cellulose, lipids, fatty acids, and flavonols (Fig. 8). Many of these enriched functions suggest a shared genetic architecture of response to abiotic and biotic stress, mediated through the action of heme oxygenases such as the diverse and biochemically active family of plant cytochrome P450 monooxygenases (450s)^65^. In plants, P450s are known to be key components of biosynthesis, metabolism, and signaling, involving responses to osmotic and temperature stress and the production of a variety of growth and defense compounds including phenylpropanoids, fatty acids, and hormones such as auxin and jasmonic acid^65,66^. Among the candidate genes repeatedly identified by all four selection scan methods, several are involved in phenylpropanoid biosynthesis (orthologs of *Arabidopsis thaliana* MYB4, CHALCONE SYNTHASE 4 (CHS4), a UDP-glycosyltransferase (UGT), and TRANSPARENT TESTA 16 (TT16) (Table 3). Phenylpropanoids are physiologically and ecologically important compounds that include lignins, anthocyanins, and condensed tannins (proanthocyanidins). Many of these compounds are known to be inducible in response to light and osmotic stress, and are also active as plant defenses against fungi, insects, and other natural enemies^67–69^. For example, MYB4 and CHS4 are both known to be responsive in *Populus* to infection by the fungal rust *Melampsora medusae*, a common leaf pathogen of poplars^70^. MYB4 expression is also responsive to jasmonic and salicylic acid, and UV-B light treatments in *Arabidopsis thaliana*^71^, while in *Populus*, CHS4 expression is induced by wounding^72,73^. A recent study of selection in 25 genes in ten steps of the condensed tannin synthesis pathway that included six CHS genes in *Populus* found elevated levels of nonsynonymous divergence at CHS genes^74^. The transcription factor PtMYB186 presents another intriguing example suggesting the action of local selection on genes with dual roles in both growth and defense (Table 3). This homolog of AtMYB206 is associated with trichome initiation – an established defense against plant herbivores; however, over-expression of PtMYB186 in transgenic poplar results in pleiotropic effects on plant growth rate, even in the absence of herbivory^75^. Overall, the functional enrichment we observed among outlier genomic regions implicates pathways, individual genes, and functional domains associated with responses to environment and plant defense, suggesting the genetic architecture of locally adaptive SGV in *Populus* may have a large pleiotropic component, shared between both abiotic and biotic stress responses.

The landscape of natural genetic variation involved in adaptation has become the increasing focus of attention of breeders, conservation biologists, and natural resource managers alike, who are facing the uncertain question of how existing populations are likely to respond to the novel pressures of global change, and where adaptive variation will be most needed in the future. Climate change conservation and mitigation strategies such as restoration or assisted gene flow^76^ increasingly require spatially-explicit solutions that rely on identifying and preserving locally adaptive SGV in different parts of a species’ range. Thus, understanding local adaptation and the distribution of adaptive SGV in the context of range position has become prerequisite for global biodiversity conservation^15,37,77,78^. Our novel combination of landscape genomics analyses has revealed the current climatic niche is of primary importance in shaping the spatial distribution of locally adaptive SGV across the range of *Populus balsamifera*, and offers a promising general approach that could be used to prioritize regions for the conservation of unique genetic resources in other crop and wild species.

## Methods

### Biological Material

We targeted southern (rear) edge and nearby regions in the core of *Populus balsamifera*’s range for sampling, spanning from northern BC to Atlantic Canada (Fig. 1). The current sampling scheme therefore covers both the rear range edge, the central core of the range, and the climatically maritime region to the east. During winter 2013, we collected dormant stem cuttings from 508 trees from 57 populations (Fig. 1) spanning 9 Canadian Provinces and 7 US States (Longitudes 55-128^*◦*^W and Latitudes 39-60^*◦*^N; Table S1). The cuttings were flushed under greenhouse conditions and fresh foliage was used for extracting whole genomic DNA using DNeasy 96 Plant Mini Kits (Qiagen, Valencia, CA, USA). DNA was quantified using a fluorometric assay (Qubit BR; Invitrogen) and confirmed for high molecular weight using 1% agarose gel electrophoresis.

### Estimation of Geographic and Climatic Range Edges

We used two distance metrics to describe the position of each *P. balsamifera* population relative to the broader range of the species: geographic distance from the southern range edge, and a measure of climatic distance from the center of the range in climatic space (Supplementary Information). Both metrics were calculated from a set of *P. balsamifera* occurrences distributed throughout the range, collected from the Global Biodiversity Information Facility and US and Canadian forest inventory programs^79,80^. To reduce spatial bias in the occurrences, we thinned the occurrences in geographic space, such that occurrences were no nearer than 75 km, resulting in 401 unique occurrences throughout the range. Occurrences were thinned using the spThin^81^ package in R.

Geographic distances from the southern range edge were calculated by first fitting an alpha hull around the 401 occurrences. Alpha hulls are similar to convex hulls, but are recommended as a way to minimize effects of spatial bias and outliers when defining a species range polygon^82^. The southern range edge was defined as the southern-most line segments of the alpha hull, from which we calculated the minimum great circle distance to each *P. balsamifera* population. Next, we used Mahalanobis distance to quantify the distance of each population from the climatic centroid of the species. We used 19 bioclimatic variables (http://www.worldclim.org/bioclim), elevation, and latitude to calculate the Mahalanobis distance, with the climatic center being defined as the average value of these variables across the 401 *P. balsamifera* occurrences. Mahalanobis distance is a unit-less, scale-invariant, multivariate distance metric that accounts for correlation among variables by scaling distances by their covariance (thereby ensuring correlated variables do not artificially inflate the measure of distance between locations). We used squared distances calculated with the Mahalanobis function in R^83^.

### GBS Sequencing

We used genotyping-by-sequencing (GBS)^84^ to obtain genome-wide polymorphism data for 508 trees. Genomic sequencing libraries were prepared from 100ng of genomic DNA per sample digested with EcoT22I restriction endonuclease followed by ligation of barcoded adapters of varying length from 4–8bp, following Elshire *et al*^84^. Equimolar concentrations of barcoded fragments were pooled and purified with QIAquick PCR purification kit. Purified products were amplified with 18 PCR cycles to append Illumina sequencing primers, cleaned again using PCR purification kit, and the resulting library was screened for distribution of fragment sizes using a Bioanalyzer. These libraries were sequenced at 48plex (i.e., each library sequenced twice) using Illumina HiSeq 2500 to generate 100bp single end reads. Cornell University Institute of Genomic Diversity (Ithaca, NY) performed the library construction and sequencing steps.

We employed the Tassel GBS Pipeline^55^ to process raw sequence reads and call variants and genotypes. In order to pass the quality control, sequence reads had to have perfect barcode matches, presence of restriction site overhang and no undecipherable nucleotides (N’s). Filtered reads were trimmed to 64bp and aligned to the *P. trichocarpa* reference assembly version 3.0^85^ using the Burrows-Wheeler Aligner (BWA)^86^. Single Nucleotide Polymorphisms (SNPs) were determined based on aligned positions to the *P. trichocarpa* reference, and genotypes called under the maximum likelihood framework in Tassel^55^. SNP genotype data along with sequence quality scores were stored in Variant Call Format v4.1 (VCF) files, which were further processed with VCFTools 0.1.11^87^. SNP genotypes were filtered to retain only biallelic sites with no indels, individuals with *<*50% missingness, sites with *<*20% missingness, genotype quality (GQ) *>*90, and site depth *>*5. We further removed SNPs that showed excess heterozygosity in tests of Hardy Weinberg Equilibrium (*P <*0.001). After filtering, the final dataset consisted of 167,324 SNPs for downstream analyses. Bioinformatic analyses were performed on a local Dell PowerEdge Linux server (24 cpu, 64 Gb RAM) at the University of Vermont and on the Teton computing environment^88^ at the University of Wyoming.

### Analysis of Population Structure

We characterized the genome-wide background genetic structure within our sample using the maximum likelihood clustering algorithm ADMIXTURE^89^. We evaluated the likelihood models for presence of up to 7 Hardy-Weinberg clusters (*K*=1 through *K*=7; Fig S2). Optimal *K* was inferred as the model that minimized the error rate based on 2000 bootstrap replicates across 10X cross-validation iterations. Cluster membership barplots were visualized using Distruct v2.3 Python script^90,91^. Individuals exhibiting ancestry of 10% or more from other *Populus* sp. were removed from further analysis (N=26).

As a complementary approach to ADMIXTURE, we used Discriminant Analysis of Principal Components (DAPC)^92^ to partition individuals along the major axes of genome-wide background genetic variation. The *K*-means clustering function used in DAPC (find.clusters()) can return different results from run to run; thus, we performed 1000 iterations of the function from *K*=1 to *K*=10, retaining the best unique solution that was most frequently observed (Fig. S1). Discriminant analyses using the most commonly observed unique solution was performed using 60% of the principal components and retaining two discriminant axes.

### Identifying Genomic Regions of Local Adaptation

Genomic signals of natural selection appear as regions of elevated population differentiation across the genomic background. Quantifying background levels of differentiation is necessary for accommodating the highly variable levels of polymorphism found within the genome, and identifying SNPs potentially be under selection. In order to generate a neutral distribution from which to infer putatively locally adapted variants, we annotated the 167,324 SNPs with VCFCodingSNPs v1.5 using the *P. trichocarpa* reference genome annotation v3.0, classifying SNPs as Downstream, Upstream, 3’ UTR, 5’ UTR, Synonymous, Non-Synonymous and Intronic. SNPs *>*5kb away from an annotated gene (hereafter called intergenic) were selected as the null set of loci for neutral parameterization of local adaptation genome scans following Lotterhos & Whitlock^93^.

For selection scans, we ranked every candidate outlier using its test statistic among the empirical null distribution of intergenic SNPs (N=1,649 for range-wide and N=1,253 for range-core data sets) and determined its empirical *P*-value. This approach caps the lower end of the empirical *P*-value proportionate with the size of the null distribution. For example, the lowest possible empirical *P*-value for the range-wide data set is 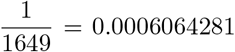 and that for the range-core data set is 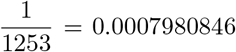. To make the inference robust, we considered an outlier a candidate if its empirical *P*-value equaled the lowest possible.

To scan for genomic regions showing elevated population differentiation, we applied the *F*-model implemented in BAYESCAN v2.1^94^ to identify SNPs under selection in range-wide as well as core groups of populations. We optimized run conditions with 20 pilot runs of 5000 MCMC sweeps each, followed by 100,000 MCMC sweeps after discarding the first 50,000 sweeps as burn-in. To minimize false positives, we used the recommendation of Lotterhos & Whitlock^93^ by setting the prior expectation of selection to one in 10,000. Candidate SNPs were then assigned empirical *P*-values based on the empirical null distribution of intergenic SNPS at an FDR of 0.01 using the empPvals() function in the R package qvalue^95^.

Additionally we estimated *X*^*T*^*X*, analogous to *F*_ST_ using the correlated allele frequencies model in BAYENV2. The maximum expected value of this statistic assuming neutrality is equal to the total number of populations being tested. To avoid spurious selection candidates, we again performed neutral parameterization to obtain empirical *P*-values for all *X*^*T*^ *X* candidate loci and only retained those with lowest empirical *P*-value.

We tested for association between SNPs and the environment using latent factor mixed models^96^ with the R package LFMM^97^. For environmental predictors, we ordinated each individual in multivariate climate space based on a PCA of the 19 bioclimatic variables and latitude. Climatic PC1 and PC2 were then used as separate predictors in association tests. Analyses were conducted on the full set of individuals with *>*90% *P. baslamifera* ancestry (N=437) and on a subset of individuals from the range core (N=336) that showed minimal background genetic structure (Fig. 1). To determine appropriate values for the number of latent factors and genomic inflation factors for our datasets, we performed a cross-validation analysis across a range of latent factors (*K*=1 to *K*=10) and at three values of the genomic inflation factor (1e-10, 1, and 1e+20). We retained parameter values that minimized the cross-validation error, and used them to construct LFMMs with the ridge penalty on a SNP-wise basis. The statistical tests of each model produced lambda-adjusted z-scores, which we used to generate empirical *P*-values based on the empirical null distribution of intergenic SNPs. We accepted only candidate SNPs with the most extreme empirical *P*-values as significant.

To allow for robust assessment of candidate outliers obtained from LFMM, we also performed gene-environment association analysis with BAYENV2^98^ on the same set of populations and climate PCs as above. First, a co-variance matrix of allele frequencies was estimated using the intergenic SNPs for range-wide and range-core data sets. Two hundred matrices were drawn (once every 5000 iterations) and the final matrix (200^th^ after 100,000 iterations) was used as a control for population relatedness in our sample. We then ran BAYENV2 on a SNP-wise basis to estimate Bayes Factors for each SNP, and ranked the outliers within the neutral distribution of intergenic SNPs to obtain empirical *P*-values. Similar to LFMM, we only retained those outliers as candidates with the lowest empirical *P*-value.

### Testing the Landscape Distribution of SGV

To investigate how locally adaptive standing genetic variation (SGV) was spatially distributed along landscape gradients of distance from the rear edge and from the climatic niche centroid, we calculated two population-level metrics of SGV based on the set of identified local adaptation outliers. We estimated a genetic index of alpha diversity (i.e., within-population variation) by calculating the allele frequency variance (*p × q*). We also calculated a genetic index of beta diversity (i.e., compositional turnover across regions) by calculating the Population Adaptive Index (PAI) *sensu* Bonin *et al*^56^. Briefly, PAI is the absolute difference between adaptive allele frequencies of a focal population and that of the entire metapopulation. It therefore represents the among-population component of SGV contributed by each focal population compared to the rest of the range. To test whether the alpha and beta diversity components of SGV varied with proximity to the rear range edge or climatic niche center, we performed a multiple linear regression between each measure of SGV and the two predictor distances. Regression models were weighted so that residual errors were inversely proportional to the sample sizes (numbers of individuals) per population (Table S1).

### Functional Enrichment

We tested for functional enrichment of genomic regions implicated in local adaptation based on the Gene Ontology (GO) terms associated with the *P. trichocarpa* reference annotation v3.0. Enrichment of GO terms associated with genes nearby to candidate SNPs was assessed based on the hypergeometric test implemented in the R package SNP2GO^99^. We tested for enrichment within a 50kb window upstream and downstream of each candidate SNP, and determined significance based on 1e+05 permutations and a false discovery rate (FDR) of 10%. Significantly enriched GO terms were then clustered for redundancy with REVIGO^100^, using the SimRel semantic similarity metric and a clustering threshold of 0.7.

## Data Archiving

Illumina sequencing data are archived under NCBI Bioproject SAMN04517823 and NCBI SRA Accession Numbers SRX1605454-68. SNP genotyping data, climate PCA loadings, and outputs from selection scans and enrichment tests are available at https://github.com/stephenrkeller/Pbalsamifera SGV.

## Acknowledgements

We are grateful to the following individuals and agencies for assistance with germplasm collections: Andrew Elmore, George Howlett Jr., Steven Guinn, Garth Inouye, and the USDA National Forest Service. We thank members of the Keller lab for feedback on the manuscript. This study was funded by a National Science Foundation award 1461868 to S.R.K. and M.C.F.

---

**Figure S1.**
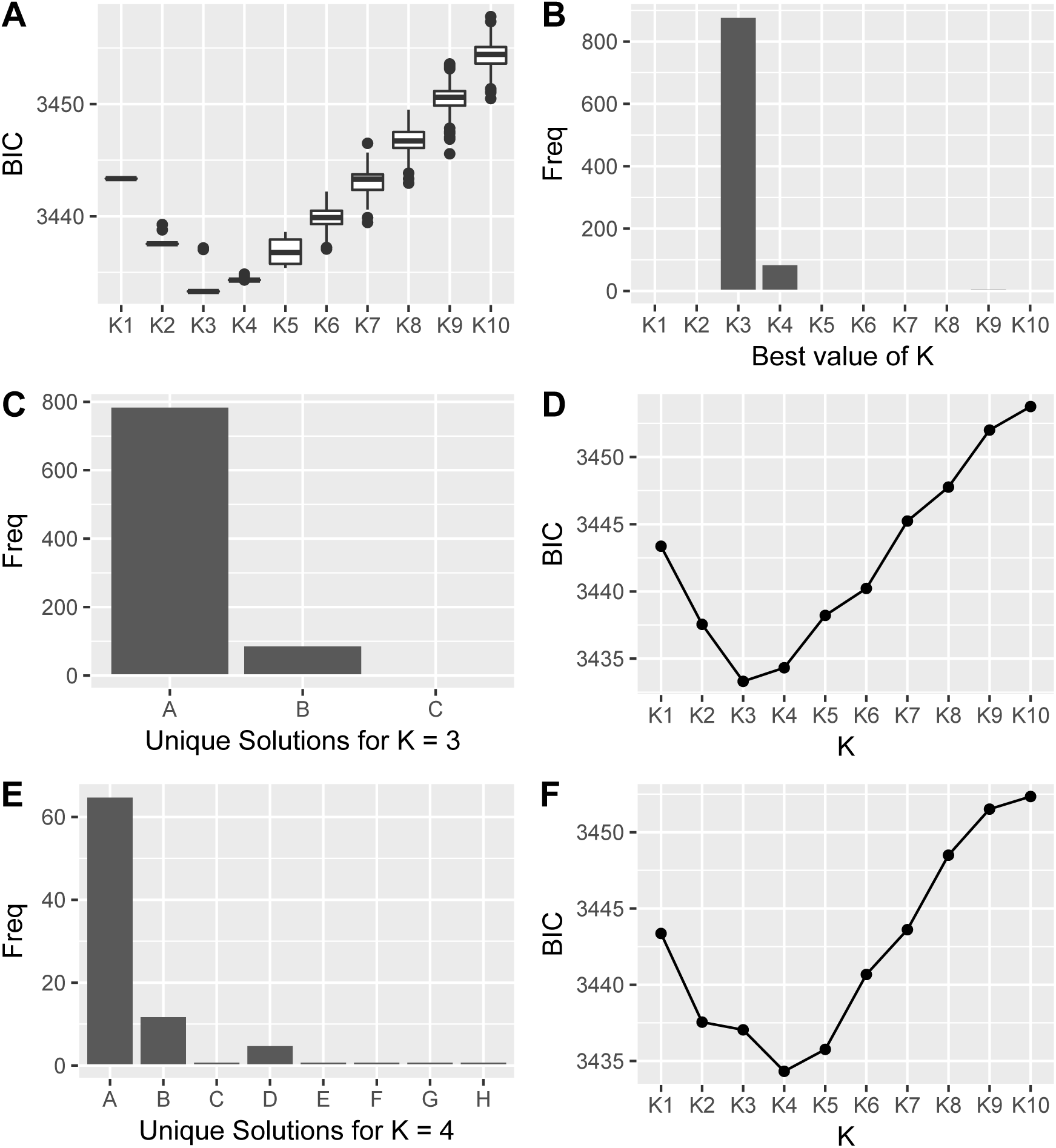
DAPC model selection output from 1000 iterations of `find.clusters` from *K*=1 to *K*=10 on the range-wide (N=437) data set (A). The best value of *K* is defined as the cluster with the lowest BIC. *K*=3 most frequently had the lowest BIC score (B). For the *K*=3 clustering results, three unique solutions were recovered (C). The BIC distribution of result A, the most common *K*=3 result (D). For the *K*=4 clustering results, more unique solutions were recovered (E). The BIC distribution of result A, the most common result at *K*=4 (F).

**Figure S2.**
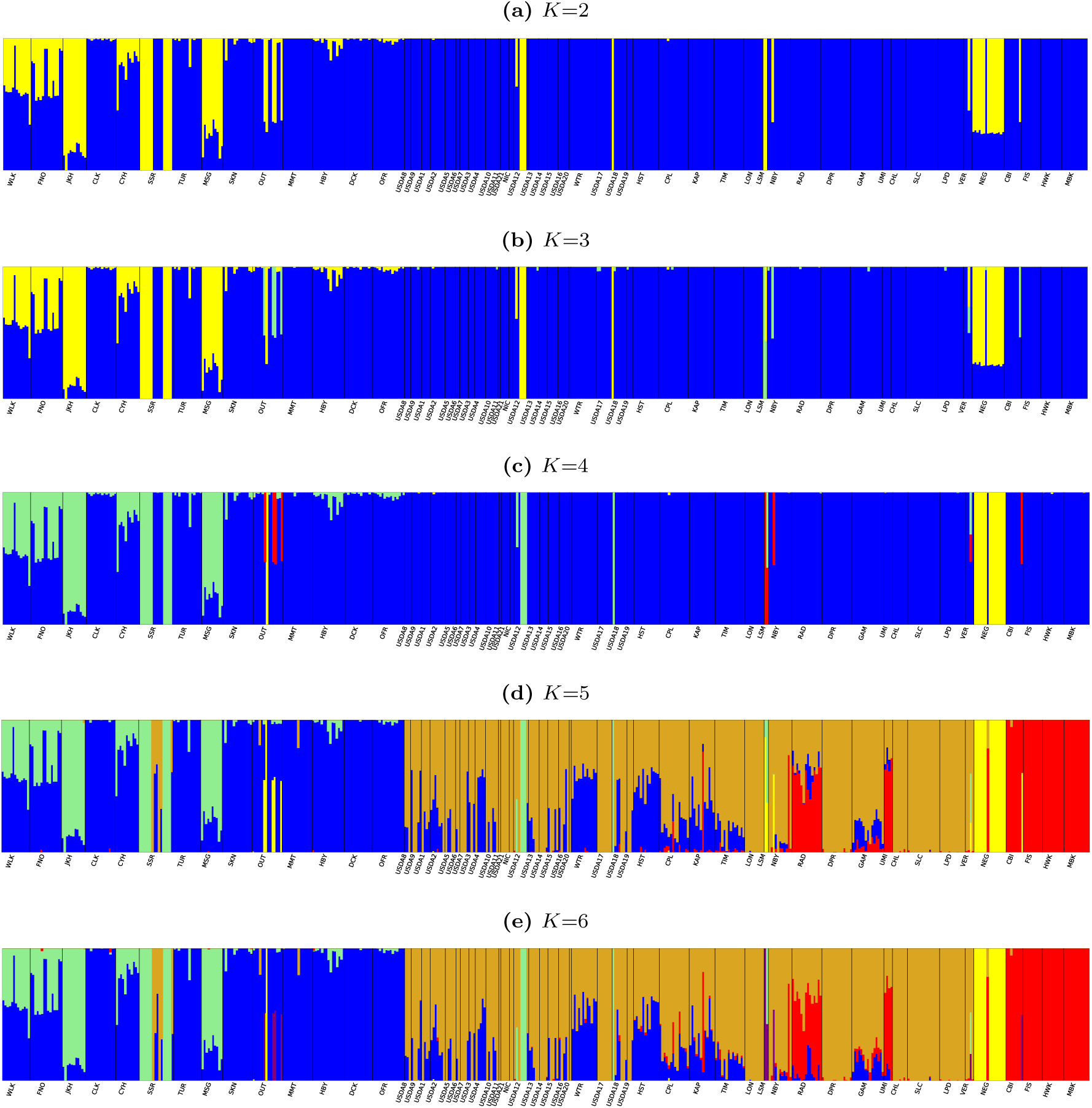
Admixture analysis of 508 individuals collected from across the range of *P. balsamifera* from *K*=2 to *K*=6.

**Table S1.**
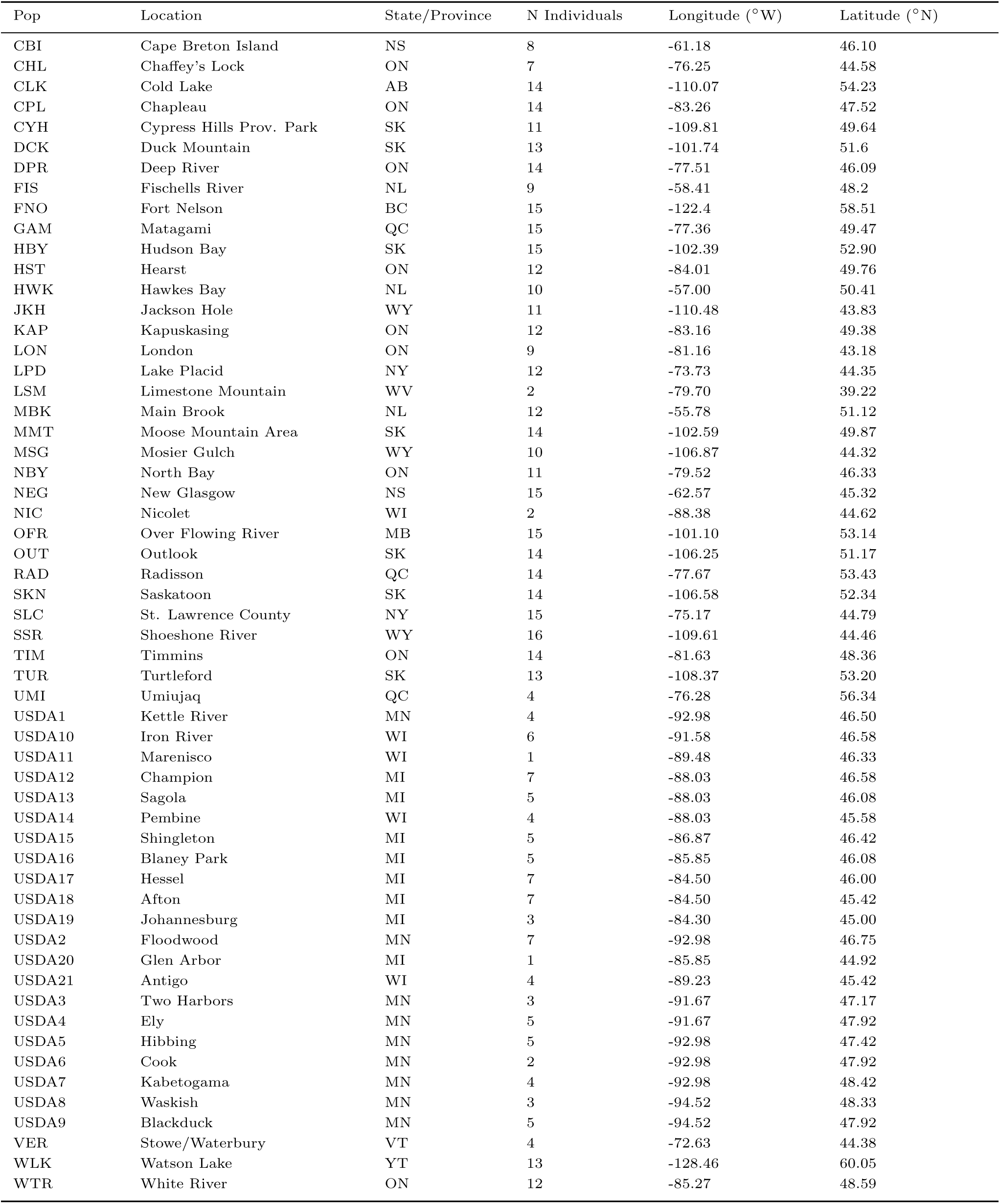
Population locality information for 508 *Populus balsamifera* individuals used in this study.

## Notes

https://github.com/stephenrkeller/Pbalsamifera_SGV

https://www.ncbi.nlm.nih.gov/bioproject/306143

